# Medium-dose irradiation impairs long-term hematopoietic stem cell functionality and hematopoietic recovery to cytotoxic stress

**DOI:** 10.1101/2024.10.29.620607

**Authors:** Qinyu Zhang, Anna Rydström, Isabel Hidalgo, Jörg Cammenga, Alexandra Rundberg Nilsson

## Abstract

Irradiation and 5-fluorouracil (5-FU) are widely utilized tools in hematopoietic research to generate myeloablation and assess blood recovery dynamics. A comprehensive understanding of their effects on the hematopoietic system is essential for optimizing therapeutic strategies, refining experimental models to modulate hematotoxicity, and interpreting research outcomes. Despite their widespread application, the long-term hematopoietic impacts of irradiation and 5-FU, particularly on hematopoietic stem cells (HSCs), remain incompletely characterized. In this study, we therefore examined the long-term effects of 2 Gy medium-dose ionizing radiation (MDIR) and 150 mg/kg 5-FU on HSCs and the hematopoietic system’s resilience to subsequent cytotoxic stress in mice. Our findings demonstrate that MDIR, but not 5-FU, induces sustained impairments in HSC function and results in the selective depletion of MHC class II^−^ HSCs – a subset characterized by high self-renewal potential. Furthermore, MDIR significantly compromised hematopoietic recovery following a subsequent 5-FU challenge, as evidenced by notably reduced platelet and red blood cell (RBC) counts during the critical recovery phase. These findings highlight the distinct and persistent impacts of MDIR and 5-FU on HSCs and hematopoietic function, revealing crucial differences in their mechanisms of action and long-term consequences on the hematopoietic system.

## Introduction

Radiation therapy and chemotherapy are foundational pillars in the treatment of cancer, commonly used as standalone interventions or in conjunction with surgery. Additionally, a combination of these modalities, known as chemo-radiation, can be utilized to maximize therapeutic efficacy. Radiation therapy employs high-energy radiation to induce DNA damage, primarily in rapidly proliferating tumor cells, leading to cell death. Chemotherapy, including agents such as 5-fluorouracil (5-FU), is administered systemically and disrupts DNA and RNA synthesis, thereby inhibiting the proliferation of rapidly dividing cells, including cancer cells, and inducing apoptosis. Beyond their clinical applications, both radiation and 5-FU are widely employed in experimental research, particularly for investigating hematopoiesis.

The hematopoietic system, undergoing continuous generation of blood cells, is particularly vulnerable to cytotoxic insults [1]. Hematopoietic stem cells (HSCs), which reside at the apex of the hematopoietic hierarchy, play a crucial role in regenerating the hematopoietic system following injury [2]. Disruptions to HSC function can thus profoundly affect the hematopoietic recovery response under stress. In experimental settings, 5-FU is commonly used to model hematopoietic stress by selectively targeting rapidly dividing cells, such as progenitors and hematopoietic precursor cells. This deletion prompts surviving HSCs to enter the cell cycle and replenish the hematopoietic system [3]. Conversely, lethal and sublethal irradiation has been the standard research strategy to ablate the hematopoietic system prior to transplantation, and to induce DNA damage for studying its biological effects. While both treatments serve as powerful tools in hematopoietic research, they also introduce biological changes that, if not properly understood, can confound experimental results. A thorough understanding of the effects of 5-FU and irradiation on HSCs and hematopoiesis is vital for designing robust experiments, accurately interpreting outcomes, and drawing valid conclusions. Clinically, this knowledge is essential for optimizing cancer treatment regimens and developing strategies to mitigate therapy-induced hematotoxicity.

In this study, we sought to investigate the long-term hematopoietic effects of 2 Gy medium-dose ionizing radiation (MDIR) and 150 mg/kg 5-FU administration in mice. While 2 Gy is reported to induce DNA damage and 10% lethality in humans [4], 150 mg/kg 5-FU is a commonly used dose in murine experimental settings [5, 6]. By examining hematopoietic recovery patterns, including changes in peripheral white blood cells (WBCs), platelets, red blood cells (RBCs), and bone marrow hematopoietic stem and progenitor cell (HSPC) dynamics, we aimed to delineate how these treatments influence hematopoietic resilience to subsequent cytotoxic stress. Our findings offer critical insights into the adaptive capacity of the hematopoietic system and lays the groundwork for optimizing therapeutic regimens.

## Results

### Distinct hematopoietic recovery patterns induced by MDIR and 5-FU

To examine the hematopoietic recovery dynamics following MDIR and 5-FU treatments, wild-type mice were subjected to a) 2 Gy irradiation, b) 150 mg/kg 5-FU, or c) volume-equivalent PBS control (Fig. 1A). Hematopoietic recovery was assessed by monitoring platelet, RBC, and WBC counts in peripheral blood over a three-month period (phase 1). Corroborating prior reports [5, 6], 5-FU treatment resulted in a marked decline in platelet and WBC concentration by day 6 (d6), followed by pronounced rebound peaks by d15, with levels stabilizing thereafter near baseline values (Fig. 1B). Subset analysis of WBCs revealed that eosinophils remained significantly depressed by d15 post 5-FU treatment and did not contribute to the overall WBC rebound peak (Figs. S1 and S2). Additionally, 5-FU caused a substantial reduction in RBCs that was followed by gradual recovery without a rebound peak (Fig. 1B). Conversely, MDIR led to significant reductions in platelet, WBC, and RBC counts, with recovery occurring gradually without any rebound peaks. Together, these results highlight the distinct peripheral blood recovery patterns induced by MDIR and 5-FU treatment, underscoring fundamental differences in how these two cytotoxic agents impact hematopoietic reconstitution.

**Fig. 1.**
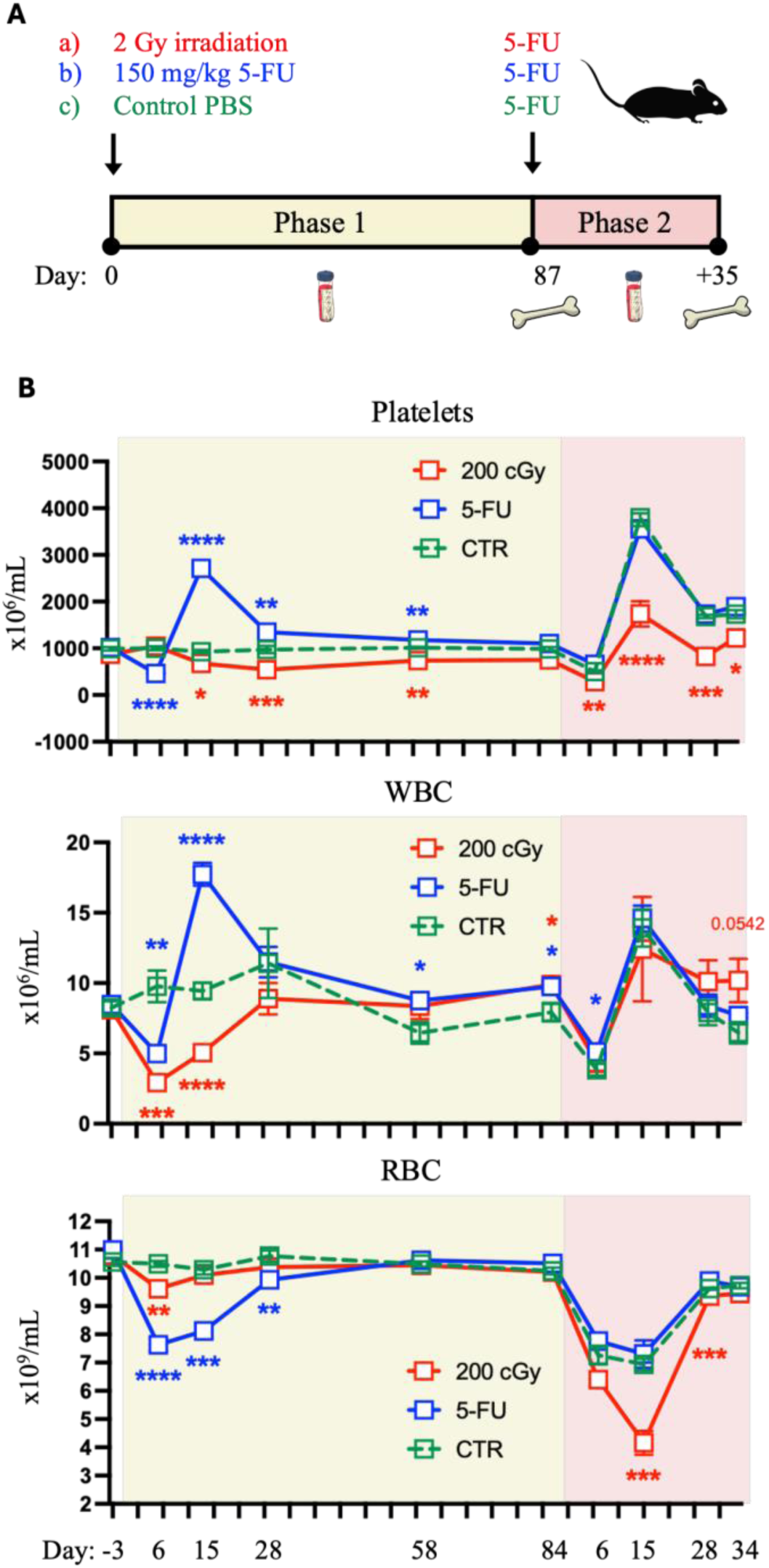
MDIR and 5-FU induce distinct peripheral blood recovery dynamics. **(A)** Experimental design. 13-week-old female C57BL/6N mice were divided into three groups: medium-dose irradiation (2 Gy), 150 mg/kg 5-FU, or control (phosphate-buffered saline, PBS). Mice were monitored for hematopoietic recovery over a 3-month period (phase 1). Following this, all groups were administered 150 mg/kg 5-FU and were observed for an additional month (phase 2). Peripheral blood counts were assessed at multiple timepoints throughout the study, with bone marrow analyses performed at phase endpoints. **(B)** Time course of peripheral blood platelet, WBC, and RBC concentrations. Sample sizes: control (n=5), 2 Gy (n=5 during phase 1, days −3 to 58; n=4 during the remainder), and 5-FU (n=5). Statistical significance is indicated for comparisons between treated groups and control. Error bars represent mean ± SEM. *p < 0.05, **p < 0.01, ***p < 0.001, ****p < 0.0001.

### MDIR results in sustained impairment of HSC functionality

To evaluate the long-term effects of MDIR and 5-FU on primitive hematopoietic populations, we assessed bone marrow HSPC subsets three months post treatments, at phase 1 endpoint. We first characterized alterations in cell counts. MDIR resulted in significant reductions in overall bone marrow cellularity (Fig, 2A) and in specific HSPC subsets, including EPCR^+^ HSCs (herein referred to as HSCs), SLAM HSCs, multipotent progenitors (MPPs), and MPPs biased towards lymphoid (MPP-Ly) and myeloid (MPP-GM) lineages (Fig 2B, gating strategies are provided in Fig. S3A and B). Megakaryocytic and erythroid-primed MPPs (MPP-MkE) were not significantly affected by MDIR. Additionally, MDIR reduced common lymphoid progenitors (CLP), pre-granulocyte/macrophage progenitors (PreGMs), pre-erythroid colony-forming units (PreCFU-Es), and megakaryocytic progenitors (MkPs), while more differentiated myeloid (GMPs) and erythroid (CFU-E/ProEry) precursor populations remained comparable to control levels. In contrast, 5-FU treatment led to a reduction in PreCFU-E precursors but did not significantly impact other analyzed HSPC subsets. To assess the functional consequences on HSCs, we isolated HSCs at the phase 1 endpoint and subjected them to *ex vivo* expansion. While HSCs from the control and 5-FU groups demonstrated similar expansion capacities, HSCs from MDIR-treated mice exhibited a pronounced reduction in *ex vivo* expansion potential (Fig. 2C, Fig. S4A). These findings reveal that MDIR, but not 5-FU, induces sustained reductions across multiple HSPC subsets and leads to a significant long-term impairment in HSC functionality.

**Fig. 2.**
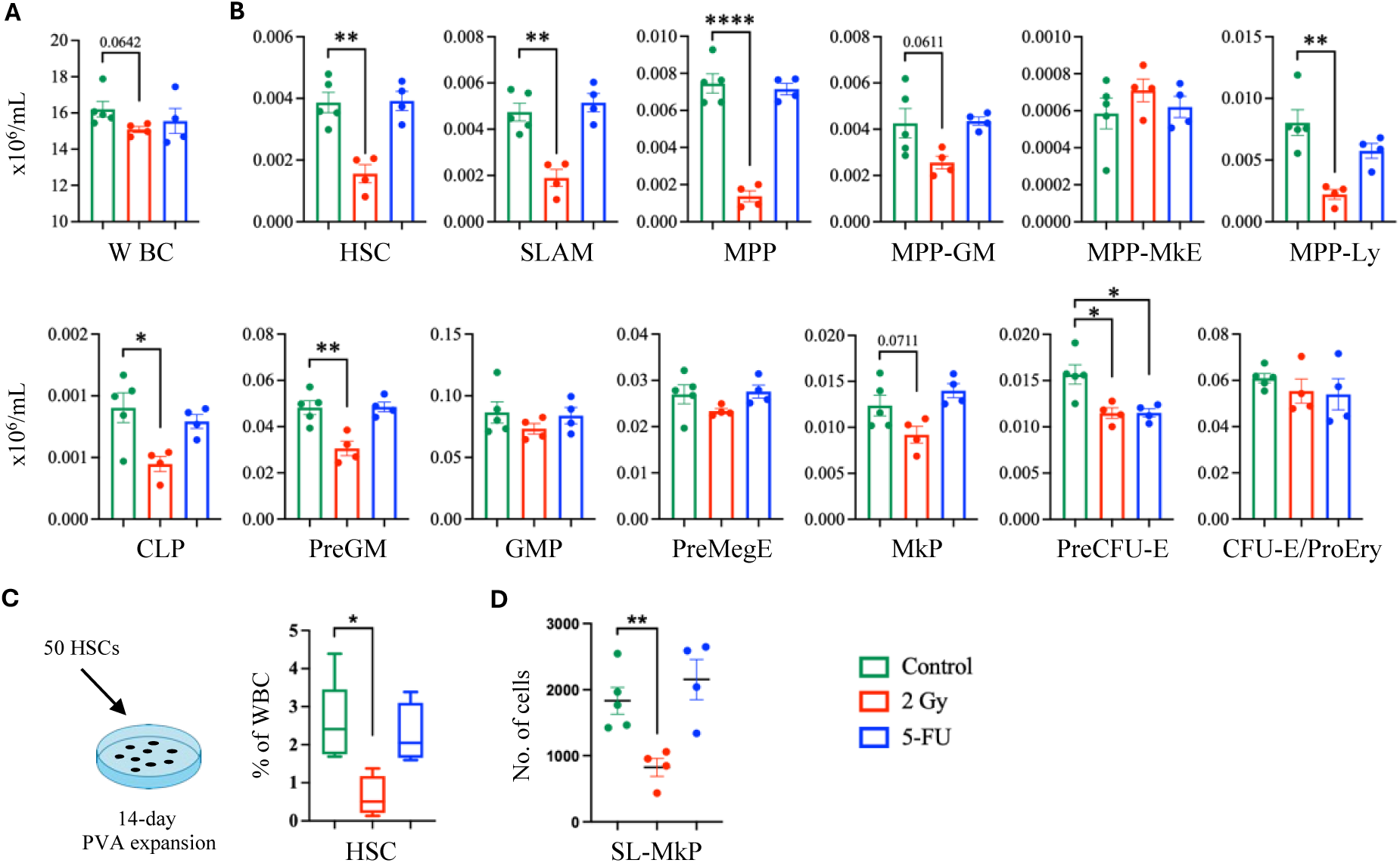
MDIR results in sustained impairments in the HSC pool. **(A)** Bone marrow WBC cellularity and **(B)** HSPC at the end of phase 1. **(C)** *Ex vivo* HSC expansion assay design (left) and frequency of HSCs after 14 days of culture (right). **(D)** Number of SL-MkPs in the bone marrow at the phase 1 endpoint. Control n=5, 2 Gy n=4, 5-FU n=4. Error bars represent mean ± SEM for panels A, B and D, and Min/Max for panel C. Statistical significance is indicated for comparisons between treated groups and control. *p < 0.05, **p < 0.01, ***p < 0.001, ****p < 0.0001.

### MDIR impairs the ability to replenish peripheral hematopoietic compartments following subsequent 5-FU treatment

To evaluate and compare the long-term impacts of MDIR and 5-FU on hematopoietic recovery ability, all groups were administered 5-FU three months after initial treatment (Fig. 1A; phase 2). While control and 5-FU only groups exhibited similar recovery patterns for RBCs, platelets, and WBCs in peripheral blood during phase 2, notable differences in RBCs and platelets were observed in the irradiated group (Fig. 1B). Irradiated mice experienced a substantially lower RBC nadir on d15 following 5-FU exposure, although all groups exhibited comparable RBC numbers at one month after the exposure (phase 2 endpoint). Additionally, irradiated mice demonstrated persistently lower platelet counts throughout phase 2, with a substantially diminished rebound peak compared to control.

The impaired platelet recovery observed during phase 2 in irradiated mice may be associated with the reduced MkP numbers observed in the bone marrow at phase 1 endpoint (Fig. 2A). Moreover, emergency platelet production can occur directly from CD41^+^ stem cell-like megakaryocyte-committed progenitors (SL-MkPs) residing within the phenotypical SLAM HSC pool [7]. Further analysis of such cells [7, 8] at phase 1 endpoint revealed that irradiated mice, but not the 5-FU group, had considerably decreased SL-MkPs numbers and frequencies (Fig. 2D, Figs. S3B and S4B). Our results demonstrate that MDIR compromises platelet and RBC production during critical recovery phases after subsequent 5-FU treatment, with reductions in MkPs and SL-MkPs likely contributing to the observed deficits in platelet output.

Finally, bone marrow analysis at phase 2 endpoint revealed no significant differences in cellularity or primitive HSPC populations between the control and 5-FU groups (Fig. 3A-C). However, the irradiated group displayed significantly reduced bone marrow cellularity, accompanied by reductions in HSCs, SLAM-HSCs, MPPs, and MPP-Ly. Notably, the observed reduction in HSC numbers was attributed to a decrease in MHC class II^−^ HSCs, whereas MHC class II^+^ HSC numbers remained unchanged (Fig. 3D and E). To evaluate functional differences between these HSC subtypes, we isolated MHC class II^+^ and MHC class II^−^ HSCs from untreated wild-type mice and subjected them to *ex vivo* expansion. MHC class II^−^ HSCs exhibited significantly greater self-renew capacity in these cultures (Fig. 3F). Collectively, these findings demonstrate that MDIR impairs the hematopoietic system’s resilience to 5-FU-mediated cytotoxic stress, with long-term alterations to both peripheral blood and primitive hematopoietic bone marrow compartments. The preferential loss of MHC class II^−^ HSCs, which exhibit enhanced self-renewal, highlights a critical and lasting impairment of hematopoietic regenerative capacity following MDIR.

**Fig. 3.**
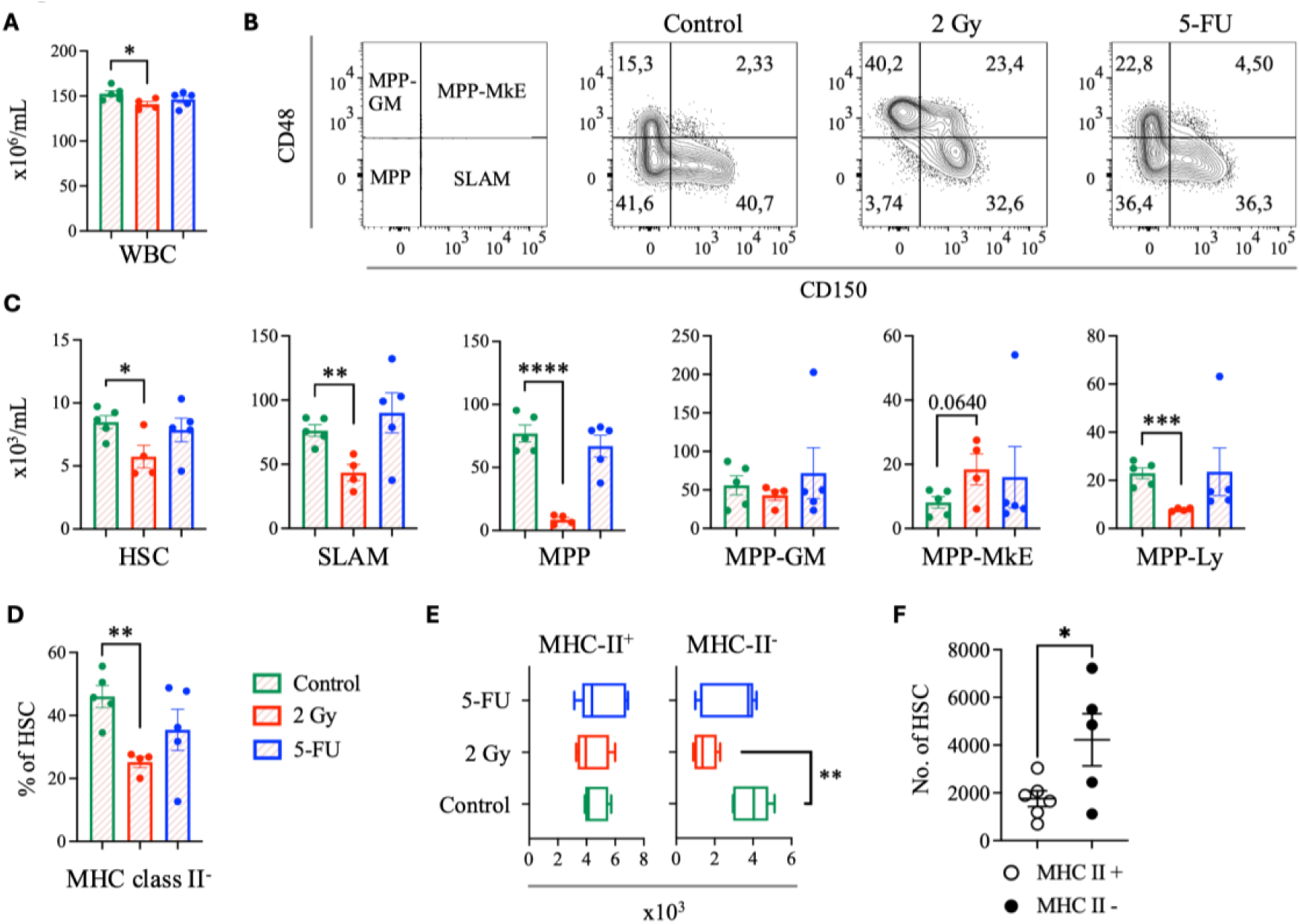
MHC class II-negative HSCs are functionally superior and are selectively depleted by MDIR. **(A)** Bone marrow WBC cellularity, **(B)** representative flow cytometry plots showing LSK populations, and **(C)** HSPC counts at the phase 2 endpoint. **(D)** Frequency of MHC class II^−^ within HSCs, and **(E)** quantification of MHC class II positive and negative HSC populations at the phase 2 endpoint. Control n=5, 2 Gy n=4, 5-FU n=5. **(F)** Number of HSCs after 13 days of *ex vivo* expansion from either MHC class II positive or negative HSCs. Control n=5, 2 Gy n=4, 5-FU n=4. Significance asterisks indicate differences in treatment groups compared to the control group. Error bars represent mean ± SEM (A, C, D and F), or Min/Max (E). *p < 0.05, **p < 0.01, ***p < 0.001, ****p < 0.0001.

## Discussion

Our study advances the understanding of the differential effects of MDIR and 5-FU on hematopoietic recovery, with a focus on the long-term consequences for HSPCs and subsequent stress responses. Through a comparative analysis of these common hematopoietic stressors, we identified unique recovery patterns and cellular responses that reveal distinct mechanisms driving hematopoietic reconstitution.

Hematopoietic recovery following 5-FU treatment featured transient reductions in peripheral blood platelet and WBC counts, followed by robust rebound peaks. RBC counts similarly declined, reaching its nadir concurrently with WBCs and platelets, but displayed delayed recovery without an overshoot. These patterns, consistent with prior studies [5, 6], suggest that 5-FU’s selective targeting of proliferating cells is followed by compensatory mechanisms to restore depleted populations. By employing control groups at each time point, we enhanced the accuracy of our findings compared to previous studies that compared the recovery levels against pre-treatment levels [5, 6], providing a clearer picture of hematopoietic recovery dynamics post-5-FU treatment. Moreover, our study uniquely characterizes WBC subset-specific responses, showing that eosinophils, unlike other subsets, fail to exhibit a compensatory overshoot. This subset-specific variability implies distinct regulatory mechanisms underlying WBC recovery and suggests that certain immune cell types may be more vulnerable to hematopoietic insults.

Prior research has shown that, unlike in the peripheral blood, 5-FU causes a transient reduction in bone marrow WBCs without a subsequent recovery peak [6, 9]. Moreover, colony-forming unit (CFU) assays have demonstrated an initial decline in CFU frequencies after 5-FU treatment, followed by an overshoot [6, 9]. However, since CFUs are primarily derived from hematopoietic progenitor cells, rather than true HSCs [10–13], CFU assays cannot reliably evaluate HSC functionality. Additionally, changes in CFU frequencies or HSC proportions may not reflect actual HSC population dynamics, as shifts in one cell type can impact the proportions of others [14]. Our results demonstrate that 5-FU does not compromise the long-term integrity or function of HSCs, with EPCR^+^ HSCs maintaining stable numbers and functionality three months post-5-FU exposure. Furthermore, exposure to a prior 5-FU dose did not modify hematopoietic recovery in response to a second 5-FU treatment, underscoring the transient impact of 5-FU on progenitor cells without affecting HSCs’ capacity for sustained hematopoietic recovery.

MDIR-induced hematopoietic suppression proved to be more persistent. MDIR triggered reductions in peripheral blood platelets, WBCs, and RBCs without rebound peaks, corroborating previous studies demonstrating the inability of MDIR to elicit a rapid compensatory response, unlike 5-FU [5, 6, 9, 15]. Previous studies have documented that MDIR results in short-term declines in bone marrow cellularity, CFUs, and SLAM HSCs [5, 6, 9, 15, 16], and long-term reductions in SLAM HSC frequencies [17]. We showed that MDIR led to lasting reductions in bone marrow cellularity and key HSPC subsets, including the more purified EPCR^+^ HSCs, which additionally exhibited impaired self-renewal capacity on a per cell basis. These enduring impacts, as opposed to the transient effects of 5-FU, indicate that radiation imposes more profound disruptions on the hematopoietic system, affecting not only cellularity but also HSC functionality and regenerative capacity.

Importantly, we also show that MDIR affects the hematopoietic recovery upon exposure to a subsequent 5-FU dose administered three months post MDIR exposure. The significantly lower platelet and RBC levels observed during the recovery phase following 5-FU in these mice, particularly the reduced platelet recovery peak, suggest a compromised emergency hematopoiesis response. This persistent impairment may arise from the sustained reduction in specific progenitor cells, such as MkPs, or platelet-biased CD41^+^ SL-MkPs that we observed in irradiated mice at the time of the 5-FU exposure, supporting previous studies indicating SL-MkPs role in emergency hematopoiesis [7]. Our data also reveal a novel vulnerability of MHC class II^−^ HSCs to MDIR, with this subpopulation showing significant reductions one month after 5-FU administration to the MDIR group, at the end of phase 2 – a finding not observed in the 5-FU only group. Given that MHC class II^−^ HSCs exhibited enhanced self-renewal capacity, their depletion may particularly impair hematopoietic recovery, underscoring the differential impacts of MDIR on distinct HSC subsets. Notably, HSCs expressing low levels of MHC class II also exhibit increased platelet potential [18], potentially further contributing to the decreased platelet output observed during phase 2 in the MDIR group. These findings have important implications for experimental models and therapeutic strategies that seek to minimize long-term hematotoxicity while enhancing HSC recovery post-radiation exposure.

In conclusion, our findings have significant implications for HSC research, where both 5-FU and irradiation are commonly used to model hematopoietic stress and investigate HSC biology. We demonstrate that MDIR and 5-FU elicit distinct effects on HSC populations and hematopoietic recovery, with MDIR inducing more enduring impairment in HSC functionality. The differential responses of HSC subpopulations, including the selective vulnerability of MHC class II^−^ HSCs to MDIR, highlight the complexity of hematopoietic responses to stress. These results emphasize the need for nuanced experimental design and interpretation, particularly in studies exploring HSC heterogeneity and regeneration. Clinically, the findings underscore the importance of careful consideration in therapeutic regimens involving radiation, where the potential for long-term hematologic complications, such as anemia or thrombocytopenia, necessitates strategies to mitigate hematopoietic impairment. Enhancing emergency hematopoiesis in irradiated patients could improve outcomes in clinical contexts, underscoring the relevance of our findings for both research and patient care.

## Material and methods

### Mice and treatments

Female C57BL/6N mice aged 12 to 13 weeks were used throughout the study. 2 Gy MDIR was delivered via exposure to a Cesium-137 source. 5-FU was administered intraperitoneally (i.p.) at a dose of 150 mg/kg body weight. Control mice received volume-equivalent doses of PBS via i.p. injections. Mice were housed at the Lund University Biomedical Center vivarium, with all procedures performed in strict accordance with ethical permits granted by the Malmö/Lund Animal Ethics Board (permit number 18055/2020).

### Peripheral blood cell monitoring and bone marrow cell analysis

Peripheral blood and bone marrow were isolated and assessed as previously described [19, 20]. Briefly, the blood was collected from the tail vein into EDTA-coated collection tubes and analyzed using a Sysmex Hemato Analyzer XN-350™ for cellularity counts. Blood samples were incubated at 37° for 25 minutes in FACS buffer (PBS containing 2% fetal bovine serum and 0.2 mM EDTA) supplemented with 1% Dextran. The WBC-rich upper phase was collected, and RBCs were lysed using RBC lysis solution (STEMCELL Technologies Inc.) at room-temperature for 3 minutes. Cells were stained with conjugated antibodies targeting CD4, CD8, NK1.1, CD19, CD11b, Gr-1, and CD45.2. Bone marrow cells were harvested from crushed tibias, femurs and iliac bones. Lineage depletion (B220, CD3, Gr-1, and Ter119) was performed prior to sorting using MACS magnetic microbead kits (Miltenyi Biotec) according to the manufacturer’s instructions. Cells were stained with conjugated antibodies for B220, CD3, Gr-1, Ter119, cKit, Sca-1, CD48, CD150, EPCR, FLT3, IL-7Ra, CD105, CD41, CD16/32, and MHC class II. Streptavidin-BV605 was used for biotin identification, and cell viability was assessed with propidium iodide (1 μg/ml; Molecular Probes). Flow cytometry analysis and cell sorting were performed on a Becton Dickinson (BD) LSR Fortessa X-20 and a BD Aria III. Cell type isolation markers and flow cytometry gating strategies are provided in the Supplementary materials.

### *Ex vivo* HSC expansion cultures

HSCs were expanded *ex vivo* following the protocol described in [21]. Briefly, 50 EPCR^+^ HSCs were sorted onto fibronectin-coated cell culture dishes containing F12 medium supplemented with 1% Insulin-transferring-selenium-ethanolamine, 1% penicillin/streptomycin/glutamine, 10 mM HEPES, 0.1% PVA (Sigma), 10 ng/mL stem cell factor and 100 ng/mL thrombopoietin. Cultures were maintained for 13 to14 days. Cellularity was quantified using a TC20 automated cell counter (Bio-Rad). Relative cell counts were assessed using flow cytometry analysis on a BD LSR Fortessa X-20, employing time-, volume, and speed-standardized settings.

## Acknowledgments

We extend our gratitude to Dr. David Bryder for his insightful feedback and valuable support throughout the course of this study. We also thank the staff at the Lund University vivarium and flow cytometry core facility for their diligent work.

## Author contributions

Conceptualization: ARN, JC

Methodology and investigation: ARN, AR, IH, QZ

Visualization: ARN

Supervision: ARN, JC

Writing: ARN

## Data and materials availability

All data needed to evaluate the conclusions in the paper are present in the paper and/or the Supplementary Materials. Key resources are listed in Table S1.

## Declaration of generative AI and AI-assisted technologies in the writing process

During the preparation of this work the author(s) used ChatGPT in order to improve language and readability. After using this tool/service, the author(s) reviewed and edited the content as needed and take(s) full responsibility for the content of the publication.

## Supplementary Figures

**Fig. S1.**
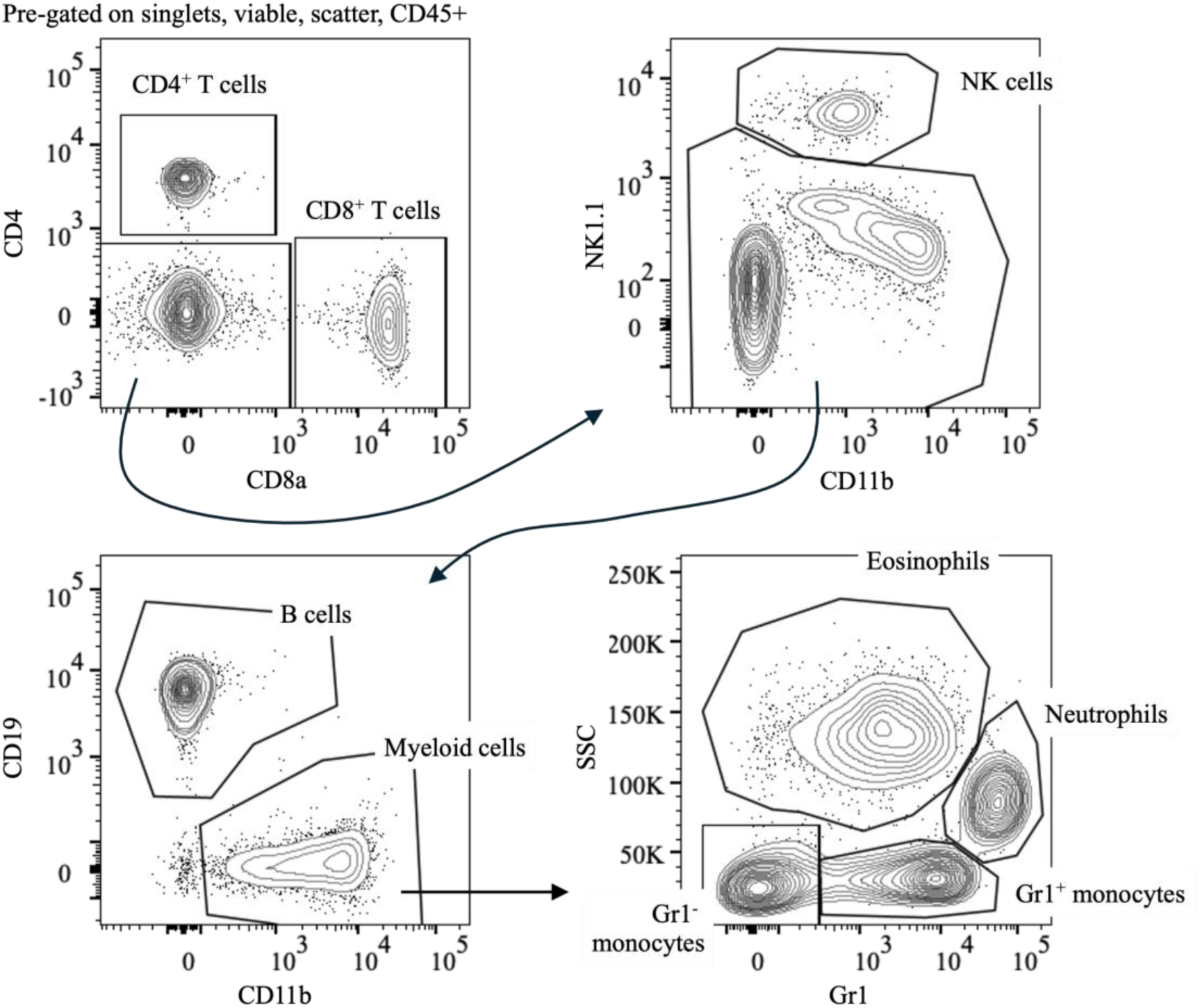
Gating strategy for peripheral blood WBC subsets. Cells were pre-gated on singlet discrimination, viability, size and complexity (FSC-A/SSC-A profiles), and CD45^+^ expression. The following subsets were identified: CD4^+^ T cells = CD4^+^CD8^−^, CD8^+^ T cells = CD8^+^CD4^−^, NK cells = NK1.1^+^CD4^−^CD8^−^, B cells = CD19^+^NK1.1^−^CD4^−^CD8^−^, Myeloid cells = CD11b^+^CD19^−^NK1.1^−^CD4^−^CD8^−^, Eosinophils = SSC^high^Gr1^low/med^CD11b^+^CD19^−^NK1.1^−^CD4^−^CD8^−^, neutrophils = SSC^med^Gr1^high^CD11b^+^CD19^−^NK1.1^−^CD4^−^CD8^−^, Gr1^+^ monocytes = SSC^low^Gr1^+^CD11b^+^CD19^−^NK1.1^−^CD4^−^CD8^−^, Gr1^−^ monocytes = SSC^low^Gr1^−^ CD11b^+^CD19^+^NK1.1^−^CD4^−^CD8^−^.

**Fig. S2.**
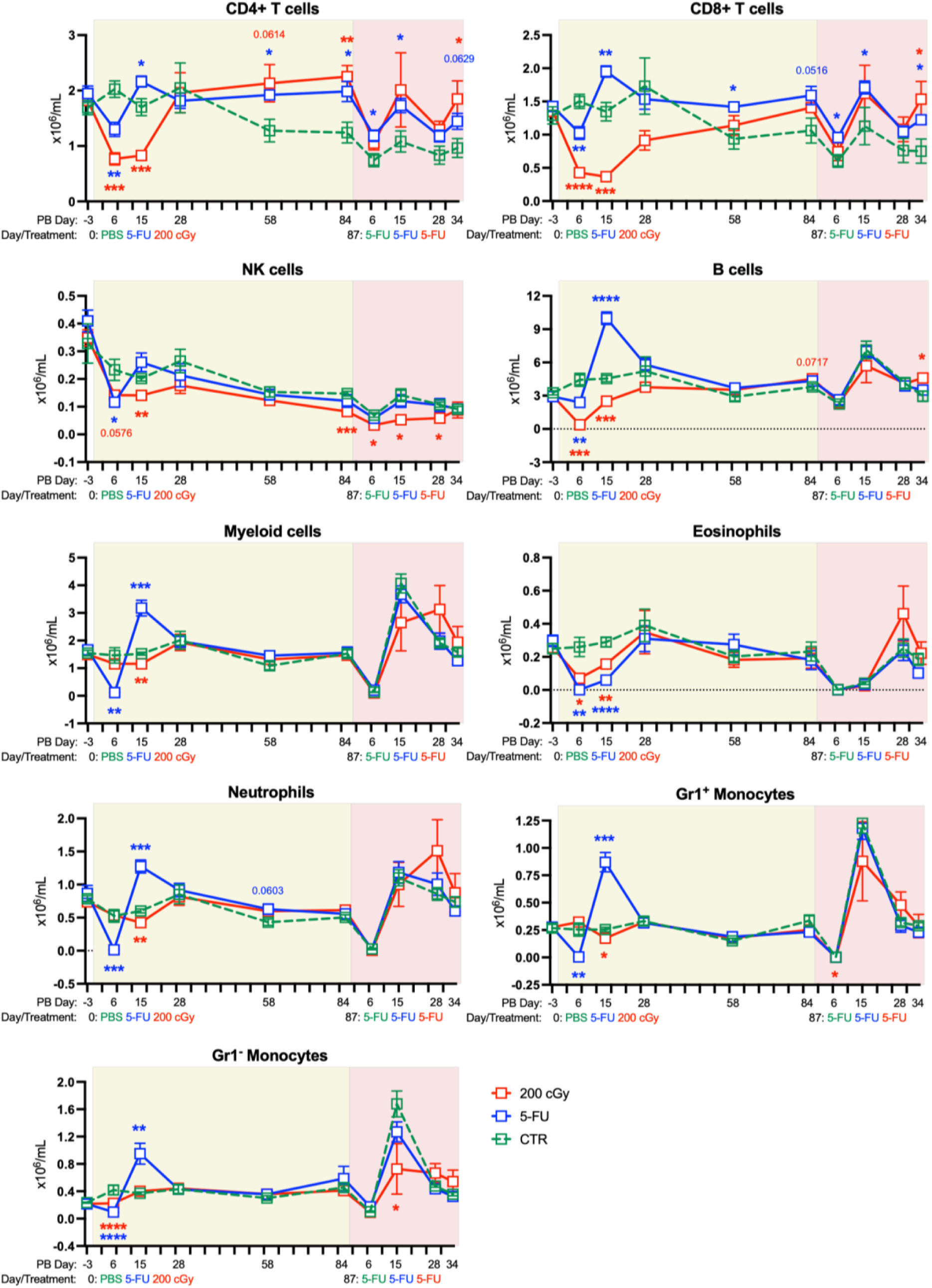
Time course of peripheral blood WBC subset concentrations. Control (n=5), 2 Gy (n=5 during phase 1, days −3 to 58; n=4 during the remainder), and 5-FU (n=5). Statistical significance is indicated for comparisons between treated groups and control. Error bars represent mean ± SEM. *p < 0.05, **p < 0.01, ***p < 0.001, ****p < 0.0001.

**Fig. S3.**
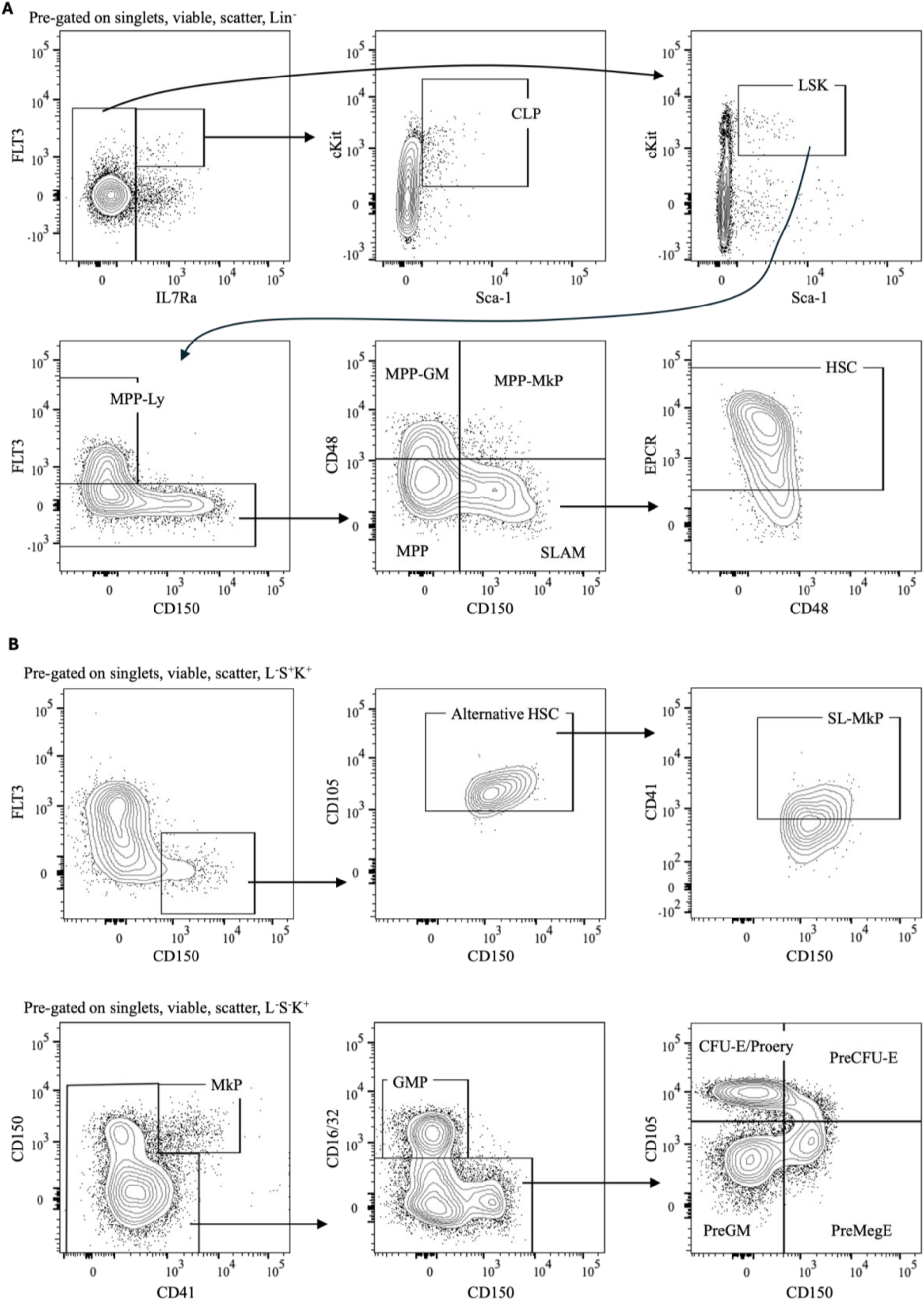
Gating strategy for bone marrow HSPC subsets. **(A)** Cells were pre-gated on singlet discrimination, viability, size and complexity (FSC-A/SSC-A profiles), and lineage (B220, CD3, Gr1 and Ter119) negativity. The following subsets were identified: CLP = FLT3^+^IL7Ra^+^Sca-1^med^cKit^med^, MPP-Ly = FLT3^+^CD150^−^IL7Ra^−^Sca-1^+^cKit^+^, MPP-MkP = CD48^+^CD150^+^FLT3^−^IL7Ra^−^Sca-1^+^cKit^+^, MPP-GM = CD48^+^CD150^−^FLT3^−^IL7Ra^−^Sca-1^+^cKit^+^, MPP = CD48^−^CD150^−^FLT3^−^IL7Ra^−^Sca-1^+^cKit^+^, SLAM = CD48^−^CD150^+^FLT3^−^ IL7Ra^−^Sca-1^+^cKit^+^, HSC = EPCR^+^CD48^−^CD150^+^FLT3^−^IL7Ra^−^Sca-1^+^cKit^+^. **(B)** Cells were pre-gated on singlet discrimination, viability, size and complexity (FSC-A/SSC-A profiles), as well as lineage (B220, CD3, Gr1 and Ter119) negativity, Sca-1^−^ and cKit^+^ (L^−^S^−^K^+^; upper panel) or L^−^S^+^K^+^ (LSK; lower panel). The following subsets were identified: SL-MkP = L^−^S^+^K^+^CD150^+^CD105^+^FLT3^−^CD41^+^, MkP = L^−^S^−^ K^+^CD150^+^CD41^+^, GMP = L^−^S^−^K^+^CD150^−^CD41^−^CD16/32^+^, PreGM = L^−^S^−^K^+^CD150^−^ CD105^−^CD41^−^CD16/32^−^, PreMegE = L^−^S^−^K^+^CD150^+^CD105^−^CD41^−^CD16/32^−^, PreCFU-E = L^−^S^−^K^+^CD150^+^CD105^+^CD41^−^CD16/32^−^, and CFU-E/ProEry = L^−^S^−^K^+^CD150^−^ CD105^+^CD41^−^CD16/32^−^.

**Fig. S4.**
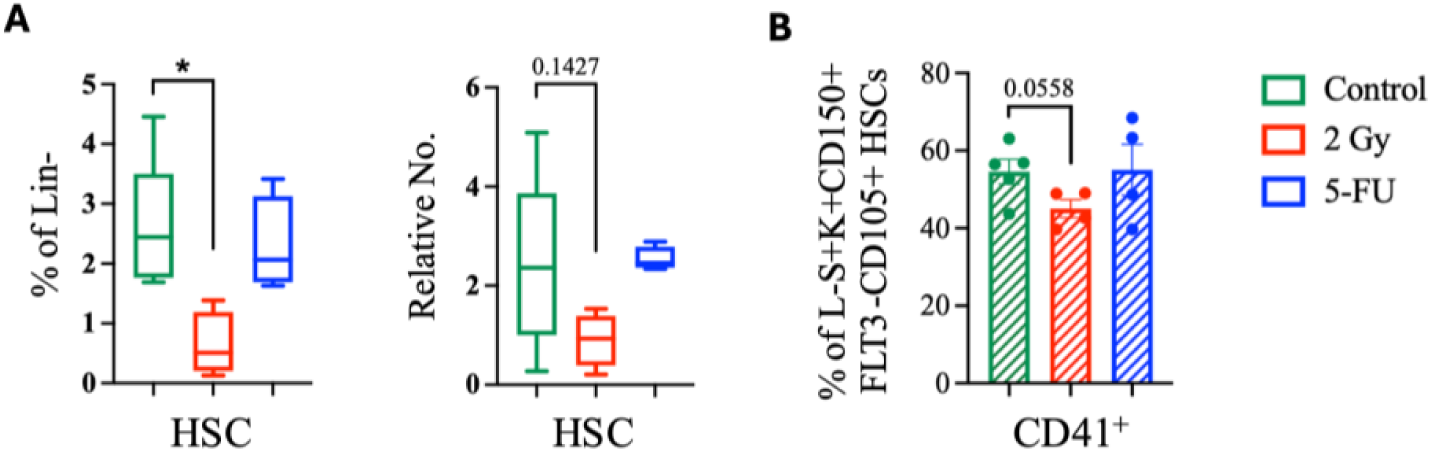
The long-term impact of MDIR and 5-FU on HSCs. **(A)** HSC frequency within the Lin^−^ compartment (left), and relative cell numbers (right) after 14 days of *ex vivo* expansion culture from HSCs isolated at phase 1 endpoint. **(B)** Frequency of CD41^+^ cells (SL-MkPs) within the alternative L^−^S^+^K^+^FLT3^−^CD150^+^CD105^+^ HSC population [8] at the end of phase 1. Control n=5, 2 Gy n=4, 5-FU n=4. Error bars represent mean ±Min/Max for panel A, and ±SEM for panel B. Statistical significance is indicated for comparisons between treated groups and control. *p < 0.05, **p < 0.01, ***p < 0.001, ****p < 0.0001.

## Supplementary tables

**Table S1.**
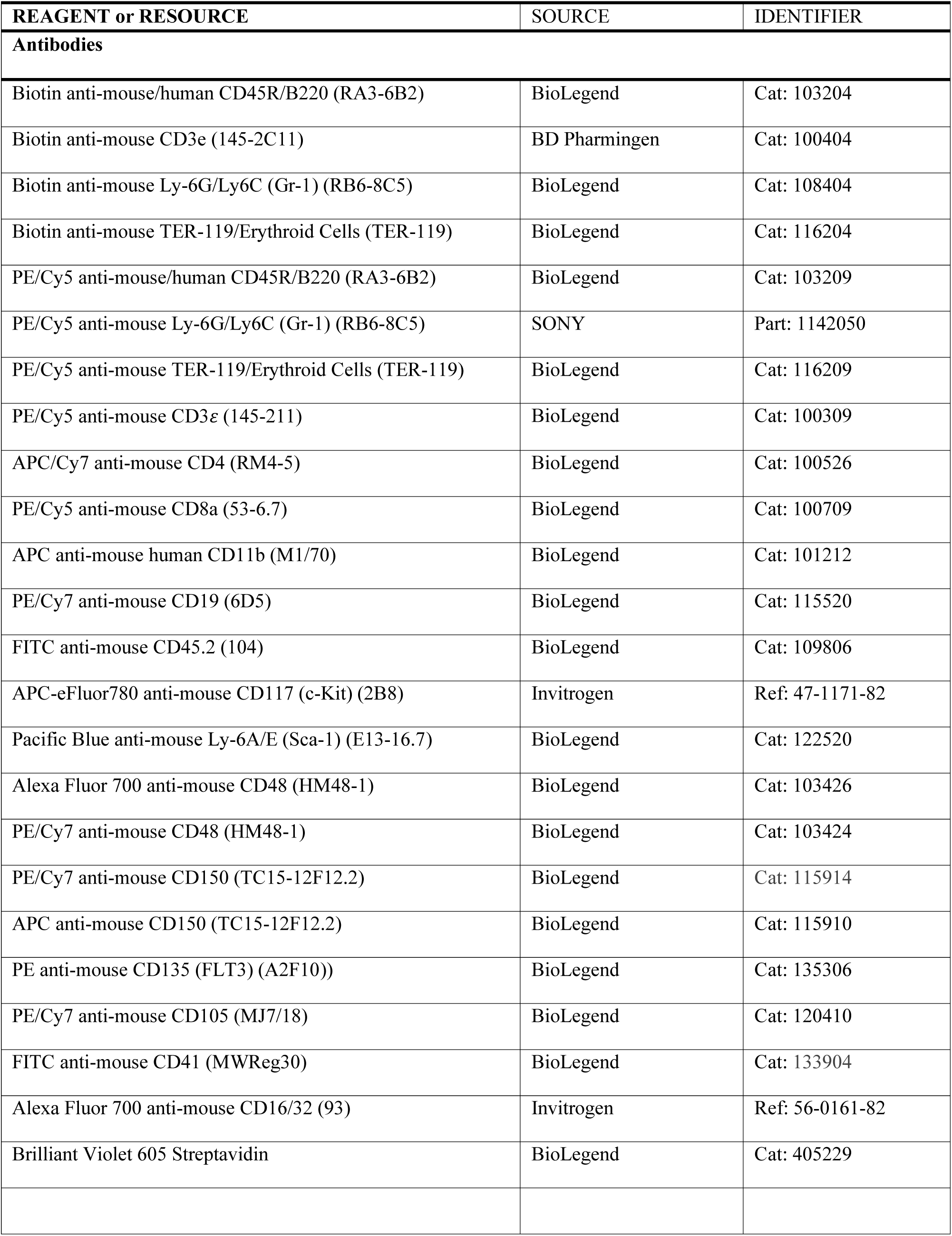
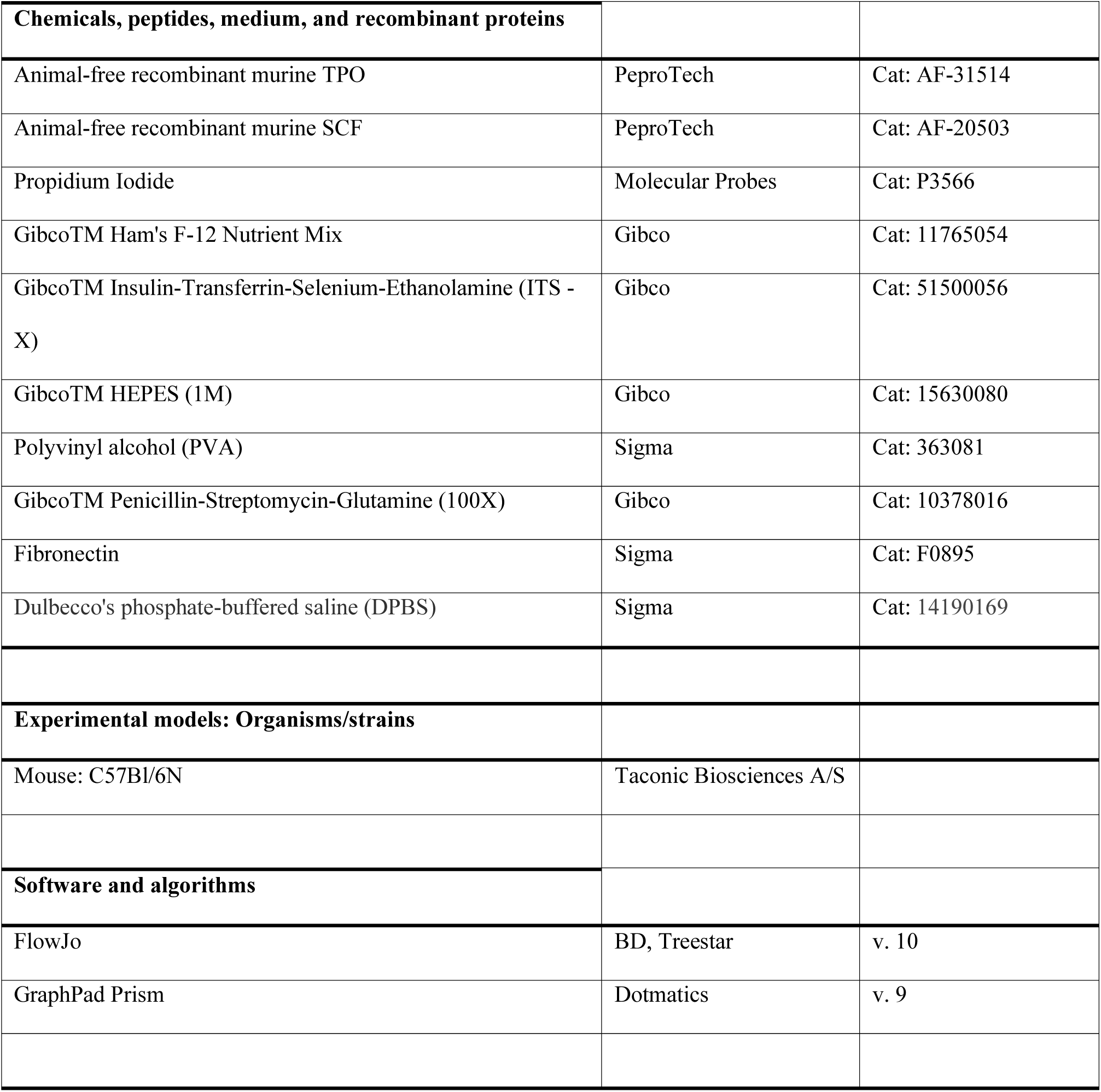
Key resource table.

## References

1. Henry, E. and M.L. Arcangeli, How Hematopoietic Stem Cells Respond to Irradiation: Similarities and Differences between Low and High Doses of Ionizing Radiations. Exp Hematol, 2021. 94: p. 11–19.

2. Pietras, E.M., et al., Functionally Distinct Subsets of Lineage-Biased Multipotent Progenitors Control Blood Production in Normal and Regenerative Conditions. Cell Stem Cell, 2015. 17(1): p. 35–46.

3. Randall, T.D. and I.L. Weissman, Phenotypic and functional changes induced at the clonal level in hematopoietic stem cells after 5-fluorouracil treatment. Blood, 1997. 89(10): p. 3596–606.

4. Squillaro, T., et al., Concise Review: The Effect of Low-Dose Ionizing Radiation on Stem Cell Biology: A Contribution to Radiation Risk. Stem Cells, 2018. 36(8): p. 1146–1153.

5. Li, X. and W.B. Slayton, Molecular mechanisms of platelet and stem cell rebound after 5-fluorouracil treatment. Exp Hematol, 2013. 41(7): p. 635–645 e3.

6. Kojima, E. and A. Tsuboi, Effects of 5-fluorouracil on hematopoietic stem cells in normal and irradiated mice. J Radiat Res, 1992. 33(3): p. 218–26.

7. Haas, S., et al., Inflammation-Induced Emergency Megakaryopoiesis Driven by Hematopoietic Stem Cell-like Megakaryocyte Progenitors. Cell Stem Cell, 2015. 17(4): p. 422–34.

8. Pronk, C.J., et al., Elucidation of the phenotypic, functional, and molecular topography of a myeloerythroid progenitor cell hierarchy. Cell Stem Cell, 2007. 1(4): p. 428–42.

9. Nielsen, O.S., H. von der Maase, and J. Overgaard, Effect of combined 5-fluorouracil and radiation on murine hematopoietic tissue. Radiother Oncol, 1988. 13(2): p. 145–52.

10. Hodgson, G.S. and T.R. Bradley, Properties of haematopoietic stem cells surviving 5-fluorouracil treatment: evidence for a pre-CFU-S cell? Nature, 1979. 281(5730): p. 381–2.

11. Van Zant, G., Studies of hematopoietic stem cells spared by 5-fluorouracil. J Exp Med, 1984. 159(3): p. 679–90.

12. Dick, J.E., et al., Introduction of a selectable gene into primitive stem cells capable of long-term reconstitution of the hemopoietic system of W/Wv mice. Cell, 1985. 42(1): p. 71–9.

13. Schofield, R., The relationship between the spleen colony-forming cell and the haemopoietic stem cell. Blood Cells, 1978. 4(1-2): p. 7–25.

14. Rundberg Nilsson, A., C.J. Pronk, and D. Bryder, Probing hematopoietic stem cell function using serial transplantation: Seeding characteristics and the impact of stem cell purification. Exp Hematol, 2015. 43(9): p. 812–7 e1.

15. Kojima, E. and A. Tsuboi, Protection of survival and hematopoiesis in irradiated mice by 5-fluorouracil. Jpn J Cancer Res, 1992. 83(7): p. 783–8.

16. Mohrin, M., et al., Hematopoietic stem cell quiescence promotes error-prone DNA repair and mutagenesis. Cell Stem Cell, 2010. 7(2): p. 174–85.

17. Rodrigues-Moreira, S., et al., Low-Dose Irradiation Promotes Persistent Oxidative Stress and Decreases Self-Renewal in Hematopoietic Stem Cells. Cell Rep, 2017. 20(13): p. 3199–3211.

18. Li, J., et al., STAT1 is essential for HSC function and maintains MHCIIhi stem cells that resist myeloablation and neoplastic expansion. Blood, 2022. 140(14): p. 1592–1606.

19. Rundberg Nilsson, A.J.S., et al., IRF1 regulates self-renewal and stress responsiveness to support hematopoietic stem cell maintenance. Sci Adv, 2023. 9(43): p. eadg5391.

20. Rundberg Nilsson, A., et al., Temporal dynamics of TNF-mediated changes in hematopoietic stem cell function and recovery. iScience, 2023. 26(4): p. 106341.

21. Zhang, Q., et al., Ex vivo expansion potential of murine hematopoietic stem cells is a rare property only partially predicted by phenotype. Elife, 2024. 12.

